# circAβ RNA drives the formation and deposition of β-amyloid plaques in the sporadic Alzheimer’s disease

**DOI:** 10.1101/2022.08.07.503077

**Authors:** Dingding Mo

## Abstract

β-amyloid peptides (Aβ) play key causal role in Alzheimer’s disease (AD). However, the mechanisms of Aβ biogenesis in sporadic AD are still largely unknown. Moreover, current AD mouse models which overexpress mutated human APP and presenilin proteins can only mimic limited characteristics of familial AD. We recently discovered an alternative Aβ production pathway from Aβ175, an Aβ peptide containing polypeptide translated from circular circAβ-a RNA generated via backsplicing of the APP gene transcript. Here, wildtype human circAβ-a RNA was overexpressed in wildtype mouse frontal cortex. Results showed that circAβ-a overexpression drove intracellular Aβ accumulation and extensive depositions of neuronal Aβ plaques in mouse brain *in vivo*. This recapitulates the critical Aβ hallmarks of sporadic AD and represents a sporadic AD mouse model. In summary, the causal relationship between circAβ RNA overexpression and AD pathology was demonstrated. This novel AD mouse model will accelerate disease-modifying drug development of this detrimental neurodegenerative disease.

## Background

Alzheimer’s disease (AD) is the most common and detrimental neurodegenerative disease with gradual loss of brain function such as memory and cognition [1-4]. The key hallmarks of Alzheimer’s disease are the formation of extracellular β-amyloid (Aβ) plaques and intracellular neurofibrillary tangles (NFTs) in patient brains [1-9]. Although the actual role of Aβ in the pathogenesis of AD is still controversial [10, 11], extensive studies, including human genetics, have confirmed that the Aβ peptide is closely associated with the development of AD pathologies [1, 12, 13]. Recent clinical trials of anti-Aβ immunotherapy on early-stage AD patients further strengthen the significant causal role of Aβ neuronal toxicity [14].

Previous studies show that Aβ is produced from β-amyloid precursor protein (APP) through sequential proteolysis by β-and γ-secretase [13, 15]. Mutations in the protein coding sequence of APP and presenilin (PS1, PS2, the catalytic subunit of the γ-secretase complex) could accelerate the process and these mutations are associated with elevated Aβ levels and amyloid plaque deposition in the brain of familial AD patients [12, 16-33]. However, gene mutations underlying familial AD only account for less than 1–5% of all AD cases globally [1, 2], and they fail to explain the sporadic AD patient majority lacking specific mutations in the APP and presenilin genes. Moreover, β-secretase is the key enzyme that controls the rate-limiting step in the amyloidogenic pathway, although it processes only abut 10% of the total APP in neuronal cells [34]. Furthermore, when β-secretase (BACE1) and its enzyme activity are moderately increased in sporadic AD patients, APP protein and γ-secretase proteolytic activity remain unchanged [35, 36]. Taken together, these lines of evidence strongly suggest that APP full-length protein may not be the main source of Aβ peptides. This points to an, as of yet, undiscovered mechanism(s) underlying the accumulation of Aβ in brains of sporadic AD patients.

Circular RNA (circRNA) is formed during processing of primary transcripts [37, 38]. Most circular RNAs are derived from protein-coding genes, regulated in various biological processes [37]. In nerve cells of the brain, circular RNAs are particularly abundant, suggesting important neural-related functions [39]. Certain circRNAs have the potential to be translated into polypeptides [40-43]. We recently found that the APP gene can also generate a circular RNA (circAβ-a) encoding a potential precursor (Aβ-related protein, Aβ175) to an Aβ peptide, the key factor of Alzheimer’s disease. These data indicate that circAβ-a has an important pathogenic ability [44]. Since the synthesis and translation of circAβ-a does not require APP gene mutations, its neurotoxicity is a potential explanation for the development of Alzheimer’s disease in patients devoid of familiar mutations [44].

The field of Alzheimer’s disease research lacks good animal models to represent the origins and development of pathological features of sporadic Alzheimer’s disease, thus limiting the efforts to transform genetic knowledge into clinical treatment drugs. Previous animal models of familial Alzheimer’s disease relied on double-transgenic or even triple-transgenic mice that overexpress human mutated APP, PSEN, and TAU proteins to obtain amyloid plaques and tangles in the mouse brain [45-48]. The APP knock-in mouse model of the humanized Aβ sequence and Swedish mutations in exon 16, Beyreuther/Iberian and Arctic mutations in exon 17, showed classic Aβ pathology, neuroinflammation and some degree of memory impairment in an ageing-dependent manner [49]. However, all these models are far from the actual pathology of patients and there is no good clinical recapitulation of human AD pathogenesis. As described previously, extensive analysis of APP protein expression in the brains of patients showed that APP is neither mutated nor overexpressed in most cases. Therefore, the overexpression of human mutated APP protein in mouse brain to produce Aβ may cause artifacts *in vivo* [50]. Therefore, currently available mouse models have obvious shortcomings and limitations; they are unable to ensure the correct evaluation of the clinical effects of drugs in humans [50].

Overexpression of wild type human APP protein in mouse brain does not produce significant amounts of Aβ, and there is no Aβ plaque formation in these animals [45-48, 50]. In contrast, expression of human wildtype circAβ-a in the mouse brain may reflect the nature of sporadic AD as the translated product of circAβ-a itself may have the ability to produce enough Aβ and generate extracellular Aβ plaques without any mutations, thus providing a useful animal model for drug screening [44]. To achieve such a goal, I constructed an AAV9 virus harboring the circAβ-a expression cassette [44] under the human SYN1 promoter (AAV9-hSYN1-circAβ-a). After injection of the AAV9-hSYN1-circAβ-a into the frontal cortex region of mouse brain, I observed robust Aβ accumulation and plaque distributions.

## Methods

### AAV9-SYN1-circAβ-a plasmid construction, AAV9 virus preparation and injection

The circAβ-a expression cassette (with intron mediated enhancement [40]) was amplified from pCircRNA-DMo-Aβ-a [44] and inserted into an AAV2 backbone vector under the human SYN1 promoter. The construct was sequenced and transfected into HEK293 cells for AAV9 virus production (named as AAV9-SYN1-circAβ-a, or AAV9-circAβ-a,) using a standard protocol. The virus titer was calculated with qPCR (3.48 × 10^13^ GC/ml). For co-injection, 1.8 µl AAV9-SYN1-circAβ-a and 0.2 µl AAV9-SYN1-Gcamp7f (which expresses cpGFP, a green fluorescent protein (GFP) variant) were injected into layer V of the left frontal cortex region (bregma, 3.0 mm; left, 1.5 mm) in 9 months old male C57 black 6 mice. For the negative control experiments, 0.2 µl AAV9-SYN1-Gcamp7f was injected. For single virus injection, 1.8 µl AAV9-SYN1-circAβ-a was injected at the same mice brain region.

### RNA isolation and Reverse transcription-polymerase chain reaction (RT-PCR)

Total RNA isolation from the AAV9 virus injected region of the left frontal cortex was performed as previously described [40] by TRIzol LS Reagent (10296010, Invitrogen). For cDNA synthesis, 0.5 µg of total RNA was used as template for reverse transcription with the SuperScript III First-Strand Synthesis System (18080044, Invitrogen) and random hexamer primers. PCR was performed with PrimeSTAR® Max DNA Polymerase (R045B, TaKaRa) and previously established circAβ-a primers (circAβ-a-R1 and circAβ-a-F1)[44] for 40 cycles.

### Western blot

Western blot was performed as previously described [40]. Briefly, injected regions (left frontal cortex) or the control region (right frontal cortex) of mouse brains were prepared in RIAPA buffer (R0278, Sigma) with protease and phosphatase inhibitor. About 40 µg total protein of the supernatants were fractioned on 8–20% precast protein gels and transferred to 0.2 µm pore-size nitrocellulose membranes (66485, Pall). For monomer Aβ detection, the pellets of brain tissues lysed in RIAPA buffer were extracted by 70% formic acid and neutralized by 5 N NaOH in 1 M Tris solution. The resulting solutions were fractioned by 8–20% precast polyacrylamide gels and transferred to 0.2 µm pore-size PVDF membranes (ISEQ00010, Millipore). Blot membranes were heated for 5 minutes in PBS for epitope retrieving before blocking. Mouse 4G8 antibody (800708, biolegend) against human Aβ was used in Aβ immunoblotting. Mouse anti GFP-Tag mAb (AE012, ABclonal) was used for Gcamp7f detection. Mouse anti-GAPDH (AC002, ABclonal) was used as loading control. HRP Goat Anti-Mouse IgG (H+L) (AS003, ABclonal) and HRP Goat Anti-Rabbit IgG (H+L) (AS014, ABclonal) were used as secondary antibodies. Signals were developed with Super ECL Plus solution (S6009M, UElandy Inc.) and visualized with the ChemiDoc MP Imaging System (Bio-Rad).

### Immunofluorescence (IF)

Three, four or seven months after AAV9-circAβ-a injection to the left frontal cortex of 9-month-old male mice, brain slices were collected using the following procedure. Anesthetized mice were perfused with PBS and 4% paraformaldehyde (PFA). Brains were fixed by 4% PFA for 24 hours followed by 24 hours dehydration initially in 15% sucrose and another 24 hours dehydration in 30% sucrose. Sections (50 μm) of the mouse brain were sectioned in a Leica CM1950 cryostat. Brain sections on slides were treated with 70% formic acid for 4 minutes or heated in citrate buffer (10 mM trisodium citrate, 0.05% Tween 20, pH 6.0) at 95 °C for 20 minutes and thrice washed with PBS. Mouse 4G8 antibody (800708, Biolegend) against human Aβ were used in immunohistostaining incubating at 4 °C overnight, followed by incubation with Alexa Fluor® 546 goat anti-mouse IgG secondary antibody (A-11003, Invitrogen) for 1 h at room temperature. Anti-GFP (chicken) was used for cpGFP immunohistostaining. For co-staining of Aβ plaques, 1 µM thioflavin-S (in PBS) (T1892, Sigma) or 1 µM QM-FN-SO_3_ (in PBS) was added to the brain sections for a one-hour incubation at room temperature and washed with PBS four times. Brain sections were also stained with X-34 Staining solution (10 µM X-34 (SML1954, Sigma), 60% PBS (vol/vol), 40% ethanol, 20 mM NaOH) for 20 minutes at room temperature. DAPI Fluoromount-G® (0100-20, SouthernBiotech) or Fluoromount-G® (0100-01, SouthernBiotech) mounting medium was used to seal the slide coverslips. Images were generated with a ZEISS LSM 900 Confocal Microscopy System.

### Reproducibility of the data

For each condition, at least 2 or 3 mice were used in the experiments and multiple slice images were captured; representative images were shown in the manuscript.

## Results

### *In vivo* overexpression of circAβ-a RNA in the mouse brain by AAV9 virus construct injection

To establish circAβ-a overexpression in the mouse brain, I injected the AAV9 virus encoding the circAβ RNA expression cassette [44] under the neuron-specific human SYN1 promoter (AAV9-SYN1-circAβ-a, Fig. 1A), into the left frontal cortex of 9-month-old C57BL/6 mice. As an injection control, an AAV9 virus encoding GCaMP7f (AAV9-SYN1-GCaMP7f), which expresses a cpGFP protein under the SYN1 promoter was co-injected. Mice were sacrificed 1.5 months post injection, and the injected brain regions were collected. Total RNA isolation was used for RT-PCR to amplify the circAβ-a RNA with our previously established divergent primers [44] (amplicon size is 499 bp) (Fig. 1B). After agarose electrophoresis, an expected RT-PCR at the 500 bp position of the DNA ladder was seen (Fig. 1B), indicating the generation of circAβ-a RNA. Furthermore, DNA sequencing of the RT-PCR products confirmed the sequence of circAβ-a amplicon (data not shown). Thus, the circAβ-a RNA was successfully overexpressed in mouse brain *in vivo*. The results are consistent with our previously established circAβ-a RNA overexpression in HEK293 cells [44].

**Fig. 1.**
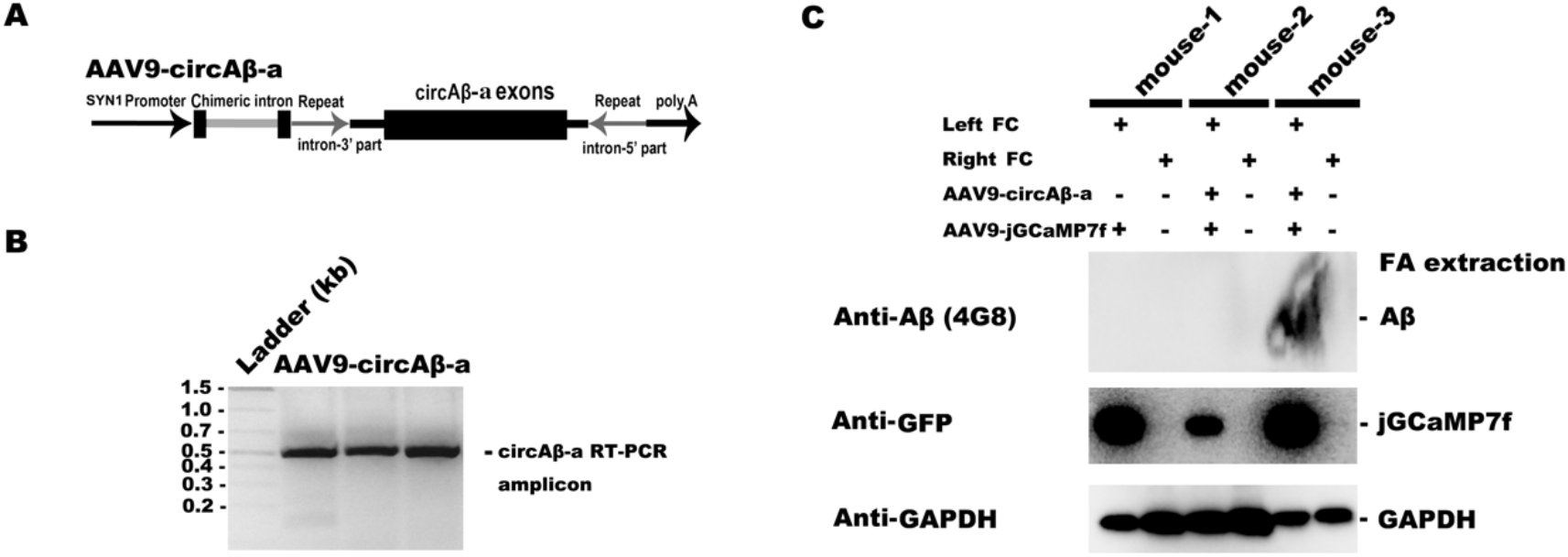
circAβ-a RNA and Aβ expression in the frontal cortex of AAV9-SYN1-circAβ-a injected mouse. A. Schematic diagram of the AAV9 virus expressing circAβ-a[44]; B. RT-PCR of circAβ-a in the frontal cortex of AAV9-SYN1-circAβ-a injected mice; the amplicon is 499 bp. C. Aβ expression in the virus injected mouse brains; Left FC, the left frontal cortex; Right FC, the right frontal cortex; Aβ were immunoblotted with 4G8 antibody; -, without; +, with.

### Aβ expression from the circAβ-a RNA in mouse brain *in vivo*

As demonstrated in HEK293 cells and in human brain samples here, circAβ-a can act as a template for Aβ175 translation and Aβ peptides are generated [44]. Total protein of the AAV virus injected region was prepared in RIPA buffer and Western blotting was performed with GAPDH and GFP antibodies for loading control. Undissolved pellets were further extracted with formic acid as detailed in Materials and Methods. These extracts were also used for the detection of Aβ by Western blot (4G8). As shown in Fig. 1C, the left frontal cortex of mouse-1 with only AAV9-SYN1-GCaMP7f injection did not reveal the corresponding signals, demonstrating that analogous injection of GFP maker virus did not affect endogenous Aβ expression from mouse APP protein. Western blot of formic acid extracted pellets from the left frontal cortex of mouse-3 revealed a heavy Aβ monomer band, yet the left frontal cortex of negative control mouse-1 failed to produce any signal (Fig. 2A). I did not detect Aβ monomer in its formic acid extracted pellet of the left frontal cortex of mouse-2, probably caused by the relatively lower injection level of AAV9-SYN1-circAβ virus and the technical difficulty of low level Aβ monomer isolation and detection (as the GFP signal is small in mouse-2) (Fig. 1C). As an endogenous negative control, the un-injected right frontal cortex does not develop any signal in the corresponding tissue areas (Fig. 1C).

**Fig. 2.**
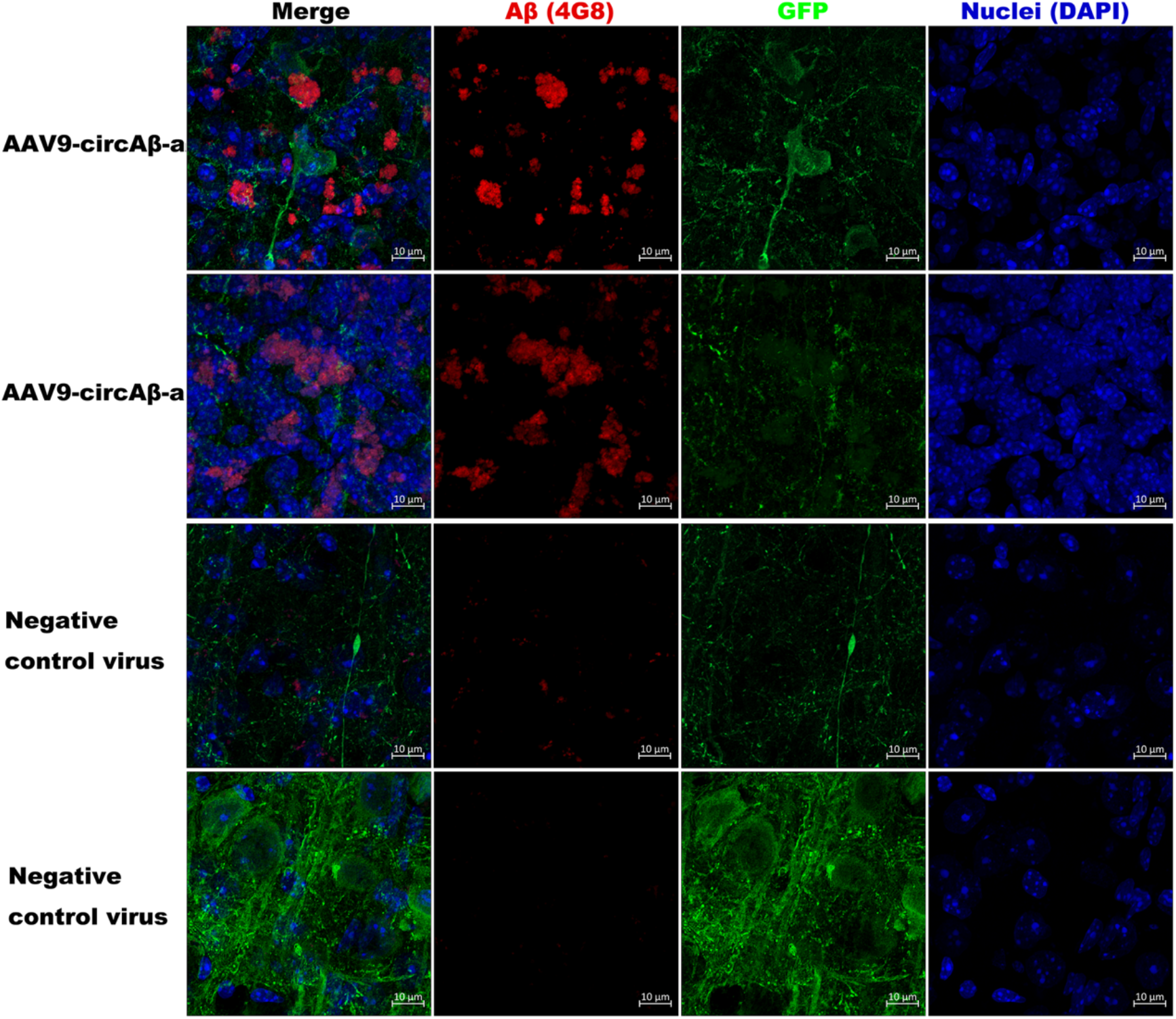
Aβ plaque depositions in the frontal cortex of AAV9-SYN1-circAβ-a injected mice. Aβ plaque immunostaining of frontal cortex after AAV9-SYN1-circAβ injection; blue, DAPI (nuclei); red, Aβ (4G8); green, GFP; Negative control virus, AAV9-SYN1-Gcamp7f.

In conclusion, Western blots showed Aβ peptides in the circAβ-a RNA overexpressed mouse brain frontal cortex tissues.

### *In vivo* Aβ plaque deposition in the circAβ-a RNA overexpressing mouse brain

Next, the question as to whether these Aβ peptides could form Aβ plaques in mouse brain *in vivo* was addressed using 4G8 antibody which can specifically identify both intracellular Aβ and extracellular Aβ plaques in mouse brain slices after 4 months of AAV9-circAβ-a virus injection. As shown in Fig. 2, the morphologies of 4G8 positive signals (red) in circAβ-a expressing tissue resembles the cotton wool Aβ plaques commonly found in AD patients [51]. A negative control showed very limited 4G8 positive signals in AAV9-SYN1-jGCaMP7f injected mouse slices. As co-injection control, the green GFP signal was prominent in both circAβ-a overexpression and negative control mice.

Furthermore, co-staining of mouse brain slices by both Aβ antibody, 4G8, and plaque-binding chemical, Thioflavin S (Thio-S), the latter binding to the β-pleated sheet conformation in amyloid plaques. In the circAβ-a expressing AAV9 virus infected mouse brain slices, I detected significant Aβ plaque signals with both with 4G8 antibody and Thio-S. Such co-localization of plaque-detecting further supports the identities of Aβ plaques (Fig.3A). Two additional Aβ plaque-detecting chemicals, namely QM-FN-SO_3_ [52] and X-34 were used to perform co-imaging of Aβ plaques with 4G8 antibody (Fig.3B, C). Once more, I obtained colocalized Aβ plaques by both 4G8 and QM-FN-SO_3_/X-34 (Fig.3B, C). Taken together, the co-staining with 3 plaque-specific chemicals and the specific Aβ antibody (4G8), solidly supported robust Aβ plaque formations and depositions in circAβ-a expressing mouse brains (Fig.2; Fig.3A, B, C). Interestingly, these positive plaques were mostly located separately from the DAPI stained nuclei (Fig.2; Fig.3A, B). Such extracellular deposits are one of the key characteristics of Aβ plaque depositions in human AD patient brains. Notably, the diameter size of these plaques was around 10 µm, relatively small compared to the average size of plaques in familial AD mouse models and AD patient brains [53, 54]. Such differences may be due to the relatively small amount of AAV virus used for infection (2 µl). Presumably, circAβ-a expressing transgenic mice will exhibit significantly larger Aβ plaques. Moreover, Aβ plaques may grow larger with the longer month after virus injection. Indeed, mice with the three different months post AAV9-circAβ-a virus injection (3, 4 and 7 months) shown the increasing sizes of Aβ plaques (Supplementary Fig. 1), demonstrating not only the reproducibility of the *in vivo* experiments but also indicating the ageing-dependence of Aβ plaques stages in the circAβ-a RNA expressing brains.

**Fig. 3.**
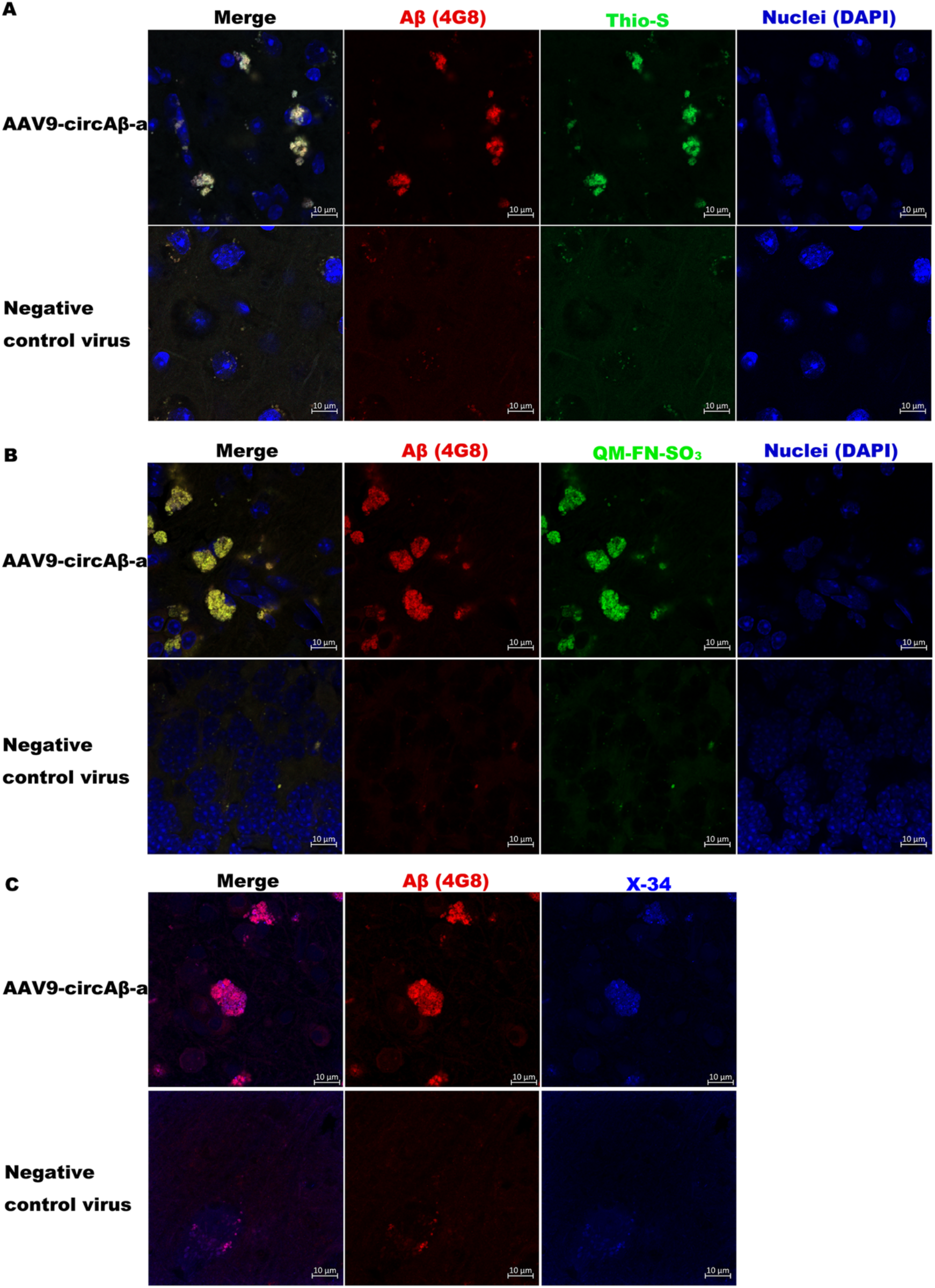
Co-staining of Aβ plaques in the circAβ-a RNA expressed mouse brain slices. **A**. blue, DAPI; green, Thioflavin S (Thio-S); red, Aβ (4G8); yellow, merged red and green. B. blue, DAPI; green, QM-FN-SO_3_; red, Aβ (4G8); yellow, merged red and green. C. red, Aβ (4G8); blue, X-34; Negative control virus, AAV9-SYN1-Gcamp7f.

### Intracellular Aβ in circAβ-a RNA overexpressing mouse brain *in vivo*

Intracellular amyloid-β not only reflects the loci of Aβ production but also plays an important role in the pathogenesis of Alzheimer’s disease, especially during the early stages [55]. Therefore, it is important to confirm intracellular Aβ expression in circAβ-a RNA overexpressing mouse brain *in vivo*. As shown in Fig. 4, the blue DAPI labels the nuclei and the red 4G8 antibody bound to all Aβ peptides; the green QM-FN-SO_3_, as shown previously, bound to aggregated Aβ plaques but not to unaggregated soluble Aβ molecules. Indeed, the arrows clearly indicates that the red 4G8 detected Aβ peptides were in the vicinity of blue nuclei, demonstrating their intracellular localization. Furthermore, they were not detected by green QM-FN-SO_3_, which detected the aggregated extracellular Aβ plaques (yellow signals in Fig. 4). In conclusion, intracellular Aβ expression occurred in the circAβ-a RNA overexpressed mouse brain *in vivo*.

**Fig. 4.**
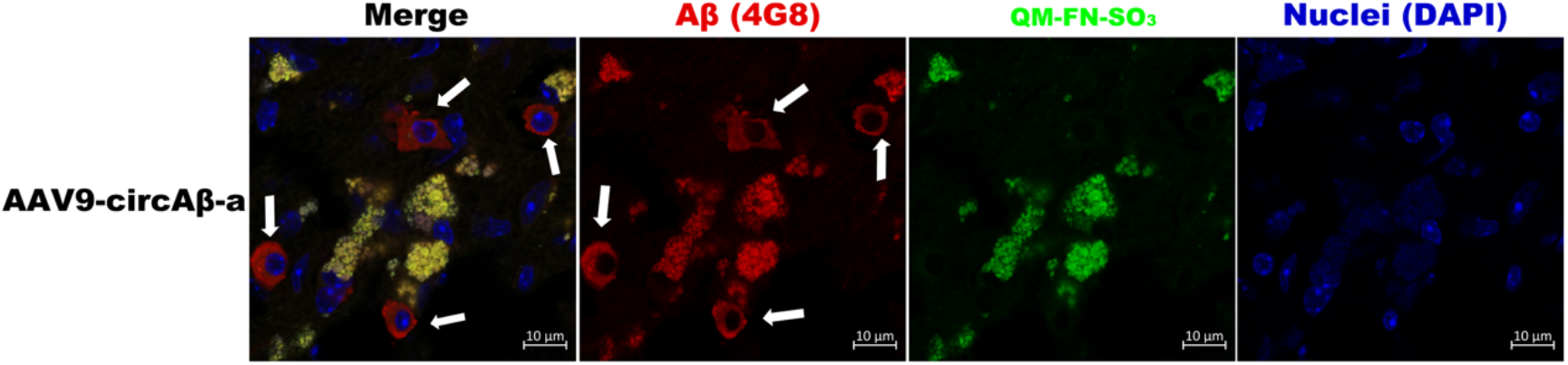
Intracellular Aβ expression in the frontal cortex of AAV9-SYN1-circAβ-a injected mouse. blue, DAPI (nuclei); green, QM-FN-SO_3_; red, Aβ (4G8); yellow, merged red and green. The white arrows indicate intracellular Aβ expression.

## Discussion

Alzheimer’s Disease is a devastating neurodegenerative disorder that threatens our ageing society. Our understanding of AD has significantly advanced since the discovery of the Aβ peptides and the derived Aβ hypothesis [1, 16, 56]. Unfortunately, disease-modifying drug development based on the Aβ hypothesis encounter challenges, reflecting the incompleteness of our current knowledge about AD molecular mechanisms [11, 56, 57]. Specifically, the main pathway for Aβ biogenesis in sporadic AD is still largely unknown and it has been assumed that the accumulation of Aβ peptides in sporadic AD brains is due to the imbalances in Aβ production or failure of Aβ clearing [11]. Such assumptions, however, fail to explain that non-mutated APP full-length proteins are not readily processed into toxic Aβ peptides [44]. Furthermore, neither APP nor PS1 proteins are mutated or upregulated in most AD patients, suggesting that there are yet undiscovered alternative Aβ biogenesis pathways in the human brain.

We recently described that circAβ-a RNA generated from APP gene transcript expresses an Aβ containing polypeptide, Aβ175; this peptide is further processed to Aβ, providing a second route for Aβ production [44]. Using multiple lines of evidence, I was able to demonstrate a robust Aβ expression in circAβ-a RNA overexpressing mouse brains *in vivo*. The circAβ-a overexpression could perfectly recreate nearly every Aβ feature of human AD brains, including the presence of intracellular Aβ and extracellular Aβ plaque depositions. Intracellular Aβ accumulation has been suggested to be an early event in AD pathogenesis, and is associated with cognitive dysfunction [55].

The circAβ-a RNA overexpressing mice produce robust intracellular Aβ, highlighting its initiating or driving role in early stages of AD (which may happen prior to the appearance of insoluble extracellular Aβ plaques). As the key hallmark of AD, robust Aβ plaque formations and depositions are observed in middle-aged mouse brains after only a three or four-month-long expression of circAβ-a RNA, further strengthening its driving pathogenetic role in AD. Interestingly, there is correlation between the Aβ plaque burden sizes and the months old of the mice and/or the expression period of the AAV-circAβ-a virus. Further study is required to elucidate the possible ageing-dependence of circAβ-a RNA translation dysregulation, presumably in a stable transgenetic mouse line.

Interestingly, the most recent cryogenic electron microscopy (cryo-EM) structures of amyloid-β 42 filaments clearly demonstrate that the so called type I filaments are predominantly found in the brains of sporadic AD patients, whereas type II filaments are majorly found in the brains of both familial AD patients as well as familial AD mouse models (AppNL-F knock-in mice) [58]. Such results support the data presented here that the morphologies of Aβ plaques observed in our circAβ-a mouse model (cotton-wool plaques) differ from those of familial AD mouse models (dense-core plaques). Cryo-EM structures of Aβ 42 filaments revealed that amyloid-β 42 (Aβ42) peptide folding differs substantially between sporadic AD and familial AD [58, 59]; the question arises as to how sequence identical peptides fold differently depending on the type of AD. It is possible that Aβ peptides are derived from different precursors. In sporadic AD, the circRNA -derived Aβ175 polypeptide is significantly shorter than the canonical APP full-length precursor protein. Consequently, it may fold into a different tertiary structure. Following secretase cleavage, the differentiating Aβ peptides remain altered, thus forming distinct Aβ filaments in sporadic AD patients compared to the full-length APP-derived Aβ peptide in familial AD.

## Conclusions

As circAβ-a RNA is well expressed in human brain [44], it is quite possible that circAβ-a RNA is the key or major pathogenic factor that expresses the majority of toxic Aβ peptides in sporadic Alzheimer’s disease (Fig. 5). In conclusion, these results significantly revise and further develop the Aβ hypothesis (Fig. 5). Specifically, these observations have elucidated the understanding of the origin of toxic Aβ forms which drive neuronal dysfunction in sporadic Alzheimer’s disease. The model described here will be invaluable to determine the factors responsible for increased Aβ expression in AD patients versus unaffected individuals. Furthermore, additional experiments targeting the production or abundance of circAβ RNA in humans by, for example, inhibiting the back-splicing reaction that generates the circular RNA from APP gene primary transcripts will be of interest. This might lead the way to next generation drugs, including antisense oligonucleotides, e.g., for reducing circAβ RNA levels, as new weapons to combat Alzheimer’s disease.

**Fig. 5.**
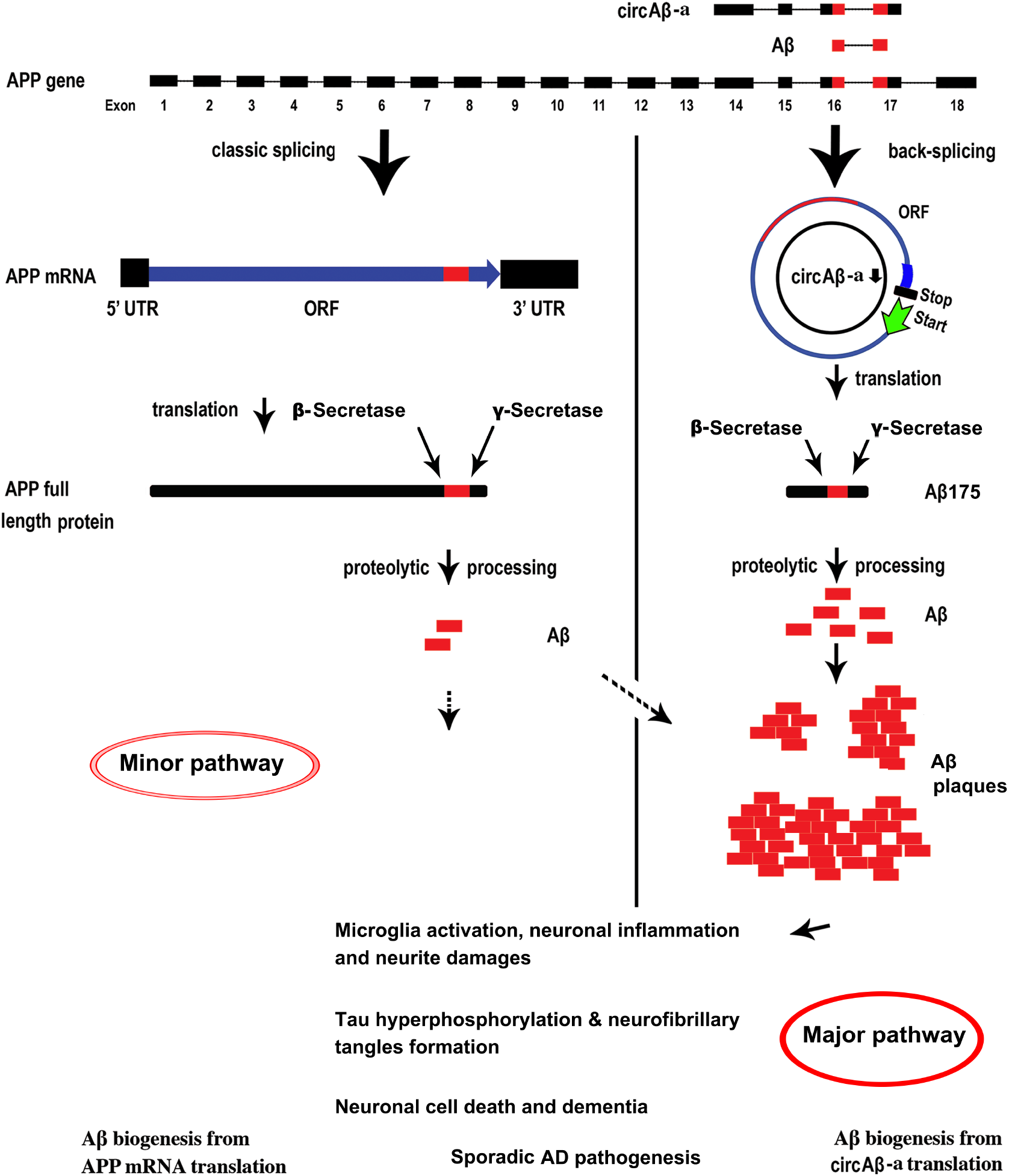
circAβ RNA drives the formation of β-amyloid plaques in sporadic Alzheimer’s disease. Proposed mechanism of Aβ biogenesis and pathogenesis in sporadic Alzheimer’s disease. Modified from the previously published original model [44].

## List of abbreviations

AAV: Adeno-associated Virus
Aβ: β-amyloid peptides
AD: Alzheimer’s disease
APP: β-amyloid precursor protein
BACE1: β-secretase 1
circRNA: circular RNA
cryo-EM: cryogenic electron microscopy
GFP: green fluorescent protein
NFTs: neurofibrillary tangles
PFA: paraformaldehyde
PS1: presenilin 1
PS2: presenilin 2
RT-PCR: Reverse transcription-polymerase chain reaction
Thio-S: Thioflavin S
X-34: 1,4-Bis(3-carboxy-4-hydroxyphenylethenyl)benzene

## Declarations

### Ethics approval and consent to participate

All the animal experiments were performed according to the corresponding laws and regulations. Detailed animal protocol was approved by the IACUC committee at Peking University Shenzhen Graduate School.

### Consent for publication

Not applicable

### Availability of data and materials

All the primary data and materials are available on request with a standard procedure.

### Authors’ contributions

D.M. designed and conceived the study. D.M. performed the experiments, analysed the results, prepared the figures, and wrote the manuscript.

## Acknowledgements

Author acknowledges Prof. Wen-Biao Gan, Dr. Wei Lei, Cong Li, Dr. Hongling Guo and Baojun Zhang for support in the research. The author also acknowledges the Imaging Core Facility of Shenzhen Bay Laboratory and Shixian Huang for assistance in slides imaging. QM-FN-SO_3_ was a gift from Prof. Zhiqian Guo. Author acknowledges Dr. Bo Zhang for sharing regents during the study. The author acknowledges Prof. Bart De Strooper, Dr. Evgenia Salta and Dr. Sriram Balusu for the supports of the study at earlier stage and comments of the manuscript. The author acknowledges Prof. Juergen Brosius for comments and revising on earlier versions of the manuscript. The author also thanks Stephanie Klco-Brosius for editorial advice.

