## Supplementary Fig. 1 for "circAβ RNA drives the formation and deposition of β-amyloid plaques in the sporadic Alzheimer’s disease"

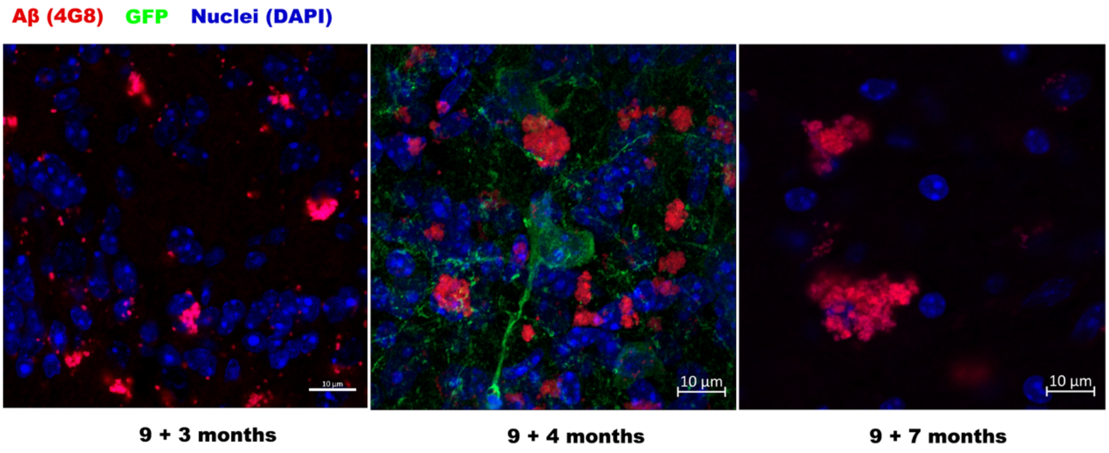


**Supplementary Fig. 1. Different sizes of Aβ plaque depositions in the mice brains of different months after AAV9- circAβ-a injection**

blue, DAPI (nuclei); red, Aβ (4G8); green, GFP. 9+3 months, 3 moths post injection of 9 months old mouse; 9+4 months, 4 moths post injection of 9 months old mouse; 9+7 months, 7 moths post injection of 9 months old mouse.
